# Perturbing whole-brain models of brain hierarchy: an application for depression following pharmacological treatment

**DOI:** 10.1101/2025.01.01.631011

**Authors:** Marcel Socoró Garrigosa, Yonatan Sanz Perl, Morten L. Kringelbach, Robin Carhart-Harris, Jakub Vohryzek, Gustavo Deco

## Abstract

Neural representation can extend beyond localised activity to encompass global patterns, where information is distributed across brain networks in a hierarchical manner. Recent research suggests that the hierarchy of causal influences shaping these patterns can serve as a signature of distinct brain states, with implications for neuropsychiatric disorders. Here, we first delve into how whole-brain models, guided by the Thermodynamics of Mind framework, can estimate the brain hierarchy of specific brain states, and how perturbations of such models can study the in-silico transitions to other states represented by static functional connectivity. We then show an application for major depressive disorder, where different brain hierarchical reconfigurations have been found following psilocybin and escitalopram treatments. We build whole-brain models of depressed patients before and after psilocybin and escitalopram interventions, and we carry a dynamic sensitivity analysis to explore the susceptibility of brain states and their drivability to healthier states. We show that susceptibility is on average reduced by escitalopram and increased by psilocybin, and that both treatments succeed in promoting healthier transitions. These results align with the post-treatment window of plasticity opened by serotonergic psychedelics, as well as with the similar clinical efficacy of both drugs observed in clinical trials.

**Graphical Abstract:** 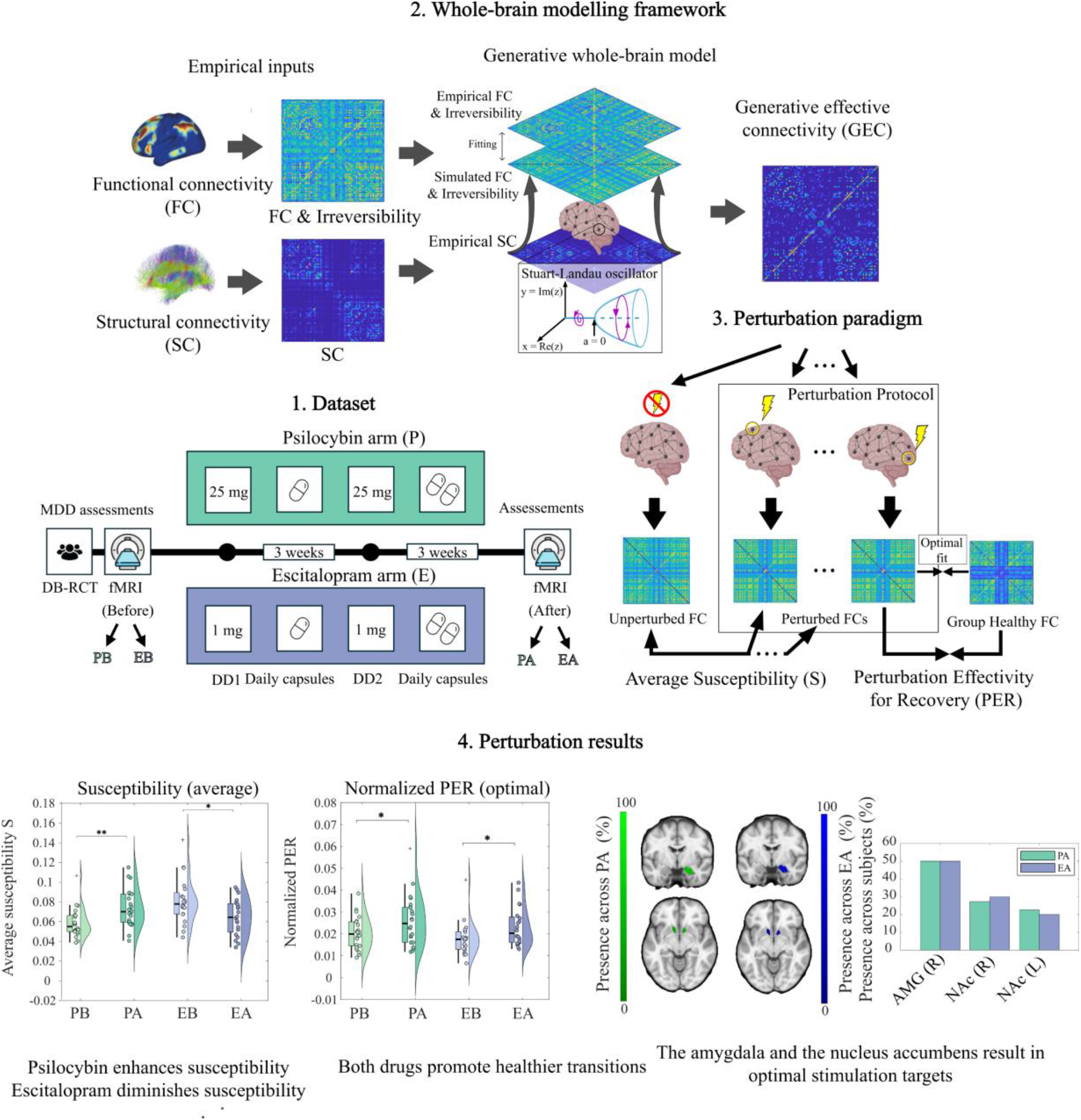

We apply whole-brain models of brain hierarchy based on the Thermodynamics of Mind framework to investigate state transitions in depression. Dynamic sensitivity analysis explores how psilocybin and escitalopram affect susceptibility and drivability to healthier states. Results show that psilocybin increases susceptibility, while escitalopram reduces it, with both enabling optimal transitions. This pipeline demonstrates the promise of in-silico approaches to inform neurostimulation protocols, potentially enhancing or complementing antidepressant therapies.

## Introduction

Understanding how the brain represents information is one of the central challenges in neuroscience. Although not entirely clear, a commonly accepted definition for *neural representation* is a “pattern of neural activity that stands for some environmental feature in the internal workings of the brain” (Vilarroya, 2017). However, a fundamental question to the notion of neural representation is at which scale does this representation occur. Historically, neural representation has been closely tied to the activity of single neurons or small clusters of neurons that encode highly specific information. This local approach stems from a wide body of research, including the discovery of place cells in the hippocampus, found to encode an organism’s spatial location (O’Keefe & Dostrovsky, 1971) or the discovery of neurons in the visual cortex which selectively respond to orientations, edges or motion (Hubel & Wiesel, 196 silico8).

More recently, modern neuroimaging techniques like functional magnetic resonance imaging (fMRI) or magnetoencephalography (MEG) have enabled researchers to move beyond the local scale and study distributed patterns of activity across the entire brain. Moreover, techniques like diffusion tractography imaging (DTI) have allowed to map the anatomical connections of the brain, giving rise to the concept of the *connectome*, defined as a “comprehensive map of neural connections in the brain on many spatial scales” (Sporns et al., 2005). Furthermore, advances in graph theory have provided mathematical tools to analyse the properties of these brain networks, giving rise to the field of *network neuroscience* (Bassett & Sporns, 2017). For instance, we now know that the brain exhibits small-worldness (Watts & Strogatz, 1998) —a balance between local specialization and global integration— and that its organization is modular (Newman, 2006).

In the context of disease, measures of functional connectivity (FC) and structural connectivity (SC) have proven to be valuable biomarkers (Greicius, 2008). Some of the examples include dementia (Rombouts et al., 2009), Alzheimer’s disease (Binnewijzend et al., 2012), posttraumatic stress disorder (Karl et al., 2006), schizophrenia (van den Heuvel & Fornito, 2014), and major depression (Greicius et al., 2007; Wang et al., 2012). However, the descriptive nature of FC and SC analyses falls short of explaining how brain structure gives rise to functional patterns.

Over a decade ago, the promising field of large-scale, whole-brain models emerged to bridge this gap (Deco & Kringelbach, 2014). Whole-brain models leverage structural connectivity data, derived from diffusion MRI, to inform the emergent large-scale brain dynamics. Based on a brain atlas, these models simulate interactions between regions by modelling their neural activity, often using mean-field approximations, such as the Wilson-Cowan model (Wilson & Cowan, 1972), or phenomenological models of coupled oscillators, like the Kuramoto and Hopf models (Cabral et al., 2017; Honey et al., 2009). In other words, they explicitly express whole-brain activity by modelling local dynamics interacting on the structural scaffold, thereby bridging the explanatory gap between structure and function of brain organization.

By fitting empirical data, these models can provide mechanistic insights into different brain states at a patient-specific level. For instance, we can optimise the anatomical connections to maximise the fit between simulated and empirical functional connectivity patterns. However, while FC connectivity is an important observable describing brain dynamics, it only captures the symmetrical relationship between different brain regions and thus fails to represent the hierarchical processing of the brain states under study. This can be overcome by using the Thermodynamics of Mind framework which considers asymmetric connectivity and thus captures the hierarchy of the causal interactions between brain regions (Deco et al., 2022; Deco et al., 2024; Kringelbach et al., 2024).

Borrowing concepts from thermodynamics, this framework acknowledges the relationship between hierarchy and non-reversibility in neural time series. Hierarchical systems reflect a structured flow of information that operates out of thermodynamic equilibrium (G-Guzmán et al., 2023; Kringelbach et al., 2024). In these systems, the asymmetries in information flow break what in physics is known as the *detailed balance*, where flux transitions between the system states disappear (Lynn et al., 2021; Sanz Perl et al., 2021). The breaking of the detailed balance establishes an arrow of time, which in the brain manifests as neural signals becoming temporally irreversible. In other words, in highly hierarchical interactions, pairwise neural time-series are causally dependent in forward time. This irreversibility or arrow of time can be estimated using temporal asymmetry by taking the difference between forward and reversal time-shifted correlations (Deco et al., 2022) (see *Materials and Methods* for details). Importantly, the irreversibility estimate is, due to its asymmetry, a robust estimate of the functional hierarchy. By incorporating the irreversibility estimates in the simulated-empirical fitting process, whole-brain models identify the causal, mechanistic generators of the functional hierarchy through the optimised anatomical connectivity (Kringelbach et al., 2024). This is known as the Generative Effective Connectivity (GEC) extending on the concept of effective connectivity (Friston, 2011).

By using generative whole-brain modelling under the Thermodynamics of Mind framework, we can create personalised models of specific brain states and uncover the underlying hierarchical causal influences between brain regions. However, the interest is often to model the transitions between brain states, for instance, from impaired to neurotypical states in neuropsychiatric disorders. A framework that allows studying plausible transitions between various brain states using whole-brain models is Dynamic Sensitivity Analysis (Vohryzek et al., 2023a), whose paradigm was introduced in a computational study on transitions between wakefulness and sleep (Deco et al., 2019). Dynamic Sensitivity Analysis consists of introducing parameter perturbations in the underlying models of local dynamics. We can interpret such perturbations as in-silico stimulations, according to the underlying model. For instance, different properties can be adjusted such as the type of stimulation (i.e., increasing excitation, modulating frequency, or adding noise, among others), the stimulation intensity, the location (i.e., local or global), and the duration (i.e., long-term, pulse-based or periodic) (Vohryzek et al., 2023a). Indeed, driving in-silico brain transitions has been shown beneficial in clinical contexts, such as in the cases of Alzheimer’s disease, psychosis and depression (Mana et al., 2024; Vohryzek et al., 2023b; Vohryzek et al., 2024a).

Major Depressive Disorder (MDD) is one of the most debilitating and problematic neuropsychiatric disorders, acutely limiting psychosocial functioning across complex and heterogeneous phenotypes (Fried et al., 2022; Malhi & Mann, 2018). Epidemiologically, it is estimated to affect 322 million people worldwide, and it is now the largest contributor to global disability and suicide deaths (World Health Organization, 2017). Moreover, it is expected to become the leading disease burden worldwide by 2030 (World Health Organization, 2008). The current gold-standard treatment is combined therapy of psychotherapy and antidepressants, which typically fall into the class of Selective Serotonin Reuptake Inhibitors (SSRIs) (Cuijpers et al., 2014). However, common antidepressants present serious adverse effects and a modest efficacy, with approximately 30% of patients showing resistance to standard treatments (McLachlan, 2018). A promising avenue in the treatment of depression lies in serotonergic psychedelics (Carhart-Harris et al., 2016; Palhano-Fontes et al., 2019). Evidence suggests that they work mainly but not only through 5-HT2A serotonin receptor (5-HT2AR) agonism, which increases serotonin release and opens a window of plasticity from which supportive psychotherapy could benefit (Carhart-Harris & Nutt, 2017).

Using generative whole-brain modelling, a recent computational study suggests that psychedelics indeed reconfigure the hierarchy of brain causal interactions very differently compared to standard SSRIs, namely by flattening, instead of increasing, the overall global directedness of the hierarchy of brain dynamics (Deco et al., 2024). This differential reconfiguration in the hierarchy suggests that psychedelic- and SSRI-treated brains should respond in different ways to stimulation protocols, which could be key to address the success of combined therapies.

Here, our aim was to study the susceptibility of brain states to perturbations and their drivability to optimal states following psilocybin and escitalopram treatments. Given the plasticity events enhanced by psychedelics, we expected brain states following psilocybin intervention to be more susceptible compared to the SSRI treatment. However, the similar efficacy in clinical trials encouraged us to hypothesise that both psilocybin and escitalopram will unlock transitions to healthier states. To answer our research question, we leveraged a double-blind phase II randomised controlled trial (Carhart-Harris et al., 2021), where resting state fMRI scans from all patients were obtained at baseline and after treatment (Daws et al., 2022). Following Deco et al. (2024), whole-brain models of each condition were built, where the Thermodynamics of Mind framework was crucial to capture the hierarchy of brain causal influences of the empirical data. Once the subject-specific models were fitted, we established a perturbation protocol following the Dynamic Sensitivity Analysis framework (Vohryzek et al., 2023a). Systematic perturbations were introduced by increasing the level of noise in the regional models, and, motivated by our hypotheses, we defined two measures to assess their effect on static functional connectivity: susceptibility, which captures the capacity of the functional pattern to change under a perturbation, and Perturbation Effectivity for Recovery (PER), which captures the similarity of the perturbed pattern and an optimal state extracted from a group of healthy participants.

## Materials and Methods

### Empirical data

The trial design and primary clinical outcomes (ClinicalTrials.gov Identifier: NCT03429075) of this study have been previously reported and registered (R. Carhart-Harris et al., 2021; Daws et al., 2022). The research was conducted at the National Institute for Health Research Imperial Clinical Research Facility under the sponsorship of Imperial College London. Ethical approval was obtained from the NHS Research and Imperial College Joint Research and Compliance Office (ID 17/LO/0389), along with approvals from the Health Research Authority and Medicines and Healthcare Products Regulatory Agency. The study was conducted in compliance with a Schedule 1 Drug Licence issued by the UK Home Office. All participants provided written informed consent and did not receive financial compensation for their involvement. This rigorous regulatory and ethical framework ensured the study’s adherence to high standards of clinical research practice and participant protection.

### Participants

To participate in the study, individuals were required to have a confirmed diagnosis of unipolar Major Depressive Disorder (MDD) from a general practitioner and a score of 16 or higher on the 21-item Hamilton Depression Rating Scale. Participants were also queried about prior use of psychedelic substances, with 31% of patients in the psilocybin group and 24% in the escitalopram group reporting previous psychedelic experience. Exclusion criteria included a personal or immediate family history of psychosis, any physician-assessed high-risk physical health condition, a history of severe suicide attempts, a positive pregnancy test, contraindications for MRI scanning, contraindications to selective serotonin reuptake inhibitors (SSRIs), or any prior use of escitalopram. Notably, treatment resistance was not considered an inclusion or exclusion criterion. All participants underwent telephone screening interviews, provided written informed consent, and completed comprehensive evaluations of their mental and physical medical histories.

### Interventions

Of the 59 patients recruited with a confirmed diagnosis of Major Depressive Disorder (MDD), 30 were randomly assigned to the psilocybin arm and 29 to the escitalopram arm using a random number generator. The final neuroimaging sample comprised 22 patients in the psilocybin arm (mean age: 44.5 years, SD: 11.0, 8 females) and 21 patients in the escitalopram arm (mean age: 40.9 years, SD: 10.1, 6 females). Prior to treatment, all participants underwent a baseline resting-state functional magnetic resonance imaging (fMRI) scan with eyes closed. On the first dosing day (DD1), participants in the psilocybin arm received 25 mg of psilocybin, while those in the escitalopram arm received 1 mg of psilocybin, a presumed negligible dose. To maintain blinding, all participants were informed they would receive psilocybin but were unaware of the specific dosage. A second dosing day (DD2) occurred three weeks later, during which patients received the same dosages as on DD1. There was no crossover of dosages between the treatment arms. From the day after DD1, patients took daily capsules for a total of 6 weeks and 1 day. In both conditions, patients ingested one capsule per day during the initial 3 weeks and increased the dosage to two capsules per day afterwards. The capsule content was either an inert placebo (microcrystalline cellulose) in the psilocybin arm or escitalopram in the escitalopram arm. In the escitalopram arm, patients received 10 mg of escitalopram for the first 3 weeks and a total of 2 × 10 mg (20 mg) thereafter.

### MRI acquisition

Brain imaging was performed on a 3T Siemens Tim Trio at Invicro. Anatomical images were acquired using the Alzheimer’s Disease Neuroimaging Initiative, Grand Opportunity (ADNI-GO56) recommended MPRAGE parameters (1-mm isotropic voxels; TR, 2,300 ms; TE, 2.98 ms; 160 sagittal slices; 256 × 256 in-plane field of view; flip angle, 9 degrees; bandwidth, 240 Hz per pixel; GRAPPA acceleration, 2). In this study, eyes-closed resting-state fMRI data were collected with T2*-weighted echo-planar images with 3-mm isotropic voxels. A 32-channel head coil was used to acquire 480 volumes in ∼10 min: TR, 1,250 ms; TE, 30 ms; 44 axial slices; flip angle, 70 degrees; bandwidth, 2,232 Hz per pixel; and GRAPPA acceleration, 2)

### Healthy target

We accessed HCP fMRI data from healthy subjects to construct a healthy target. We focused on 100 unrelated subjects from the project Young Adult, as employed in a prior study (Vohryzek et al., 2020). Participants underwent four rs-fMRI sessions of 14 min 30 s with a repetition time of 0.72 s on a 3-T Skyra scanner (Siemens). We focused on DK80 regional fMRI signals corresponding to data from the first scanning session. Acquisition and pre-processing details can be found at the HCP website (http://www.humanconnectome.org/).

### Structural connectivity template

Given that the psilocybin and escitalopram trial did not include diffusion MRI scans, we accessed a standard structural connectivity (SC) template. This connectivity was estimated using probabilistic tractography in a prior study from two dMRI projects of the Human Connectome Project (HCP) (Deco et al., 2021).

### fMRI data processing

The functional magnetic resonance imaging (fMRI) data underwent preprocessing using a custom pipeline that integrated tools from multiple neuroimaging software packages, including the FMRIB Software Library (FSL)13, Analysis of Functional NeuroImages (AFNI), FreeSurfer, and Advanced Normalisation Tools (ANTs). This preprocessing protocol, previously described in detail (Daws et al., 2022), comprised de-spiking, slice-time correction, motion correction, brain extraction, rigid-body registration of functional images to anatomical scans, nonlinear registration to a standard template, scrubbing, bandpass filtering, and nuisance regression. The fMRI DK80 parcellated time series used in the present study were extracted from this preprocessed data.

### Empirical measures

Using the DK80 parcellated time series, empirical covariances 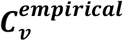 and functional connectivity (FC) matrices ***FC***^***empirical***^ were constructed by computing for each ij-th pair of regional signals (*x*_*i*_(*t*), *x*_*j*_(*t*))

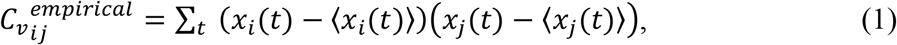

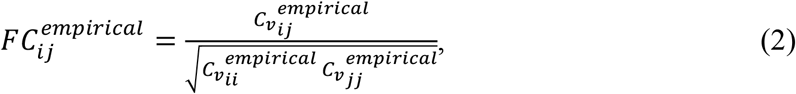

where ⟨ ⟩ indicates averaging across the time series. Similarly, we defined forward *τ* time-shifted empirical covariances ***C***_***v***_(*τ*) and forward *τ* time-shifted correlation matrices ***FC***(*τ*) as

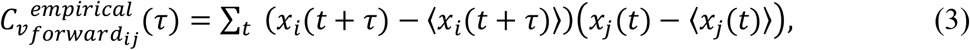

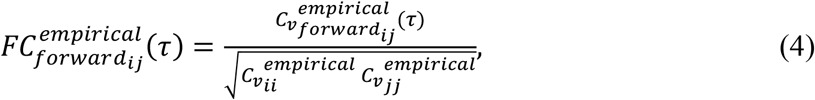

where *τ* is the shifting time, which was set to 2 times the repetition time. Note that shifting makes 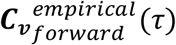 and 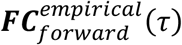 asymmetric matrices.

### Whole-brain model

We modelled local dynamics of each brain region with Stuart-Landau oscillators. This model corresponds to the normal form of a supercritical Hopf bifurcation and is widely used to investigate the transition from noisy to oscillatory dynamics (Kuznetsov, 1998). It has been used to replicate important features of brain dynamics observed in electrophysiology (Freyer et al., 2011, 2012), magnetoencephalography (Deco et al., 2017a) and fMRI (Deco et al., 2019; Kringelbach et al., 2020). Mathematically, let us consider a network of N parcellated regions. Each region j is represented as a node in the network, and the connectivity from node *k* to node *j* is captured by *C*_*jk*_ in the connectivity matrix ***C***. Whole-brain dynamics can then be modelled by N Stuart-Landau oscillators coupled through the connectivity matrix *C*.

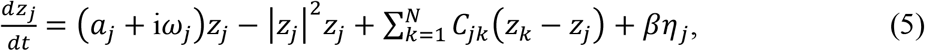

where for each region j, the complex variable *z*_*j*_ denotes the state (*z*_*j*_ = *x*_*j*_ + *iy*_j_ ), 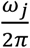 is the intrinsic frequency (in the 0.008-0.08Hz band), *βη*_*j*_ is additive uncorrelated Gaussian noise with standard deviation *β* (for all *j*), and *a*_*j*_ is the bifurcation parameter. Intrinsic frequencies were estimated from regional narrowband blood-oxygen-level-dependent BOLD signals by taking their patient-averaged peak frequency. The bifurcation parameter *a*_*j*_ places the system in a noisy regime if *a*_*j*_ < 0 and in a stable limit cycle with self-sustained oscillations if *a*_*j*_ > 0. The point *a*_*j*_ = 0 hence represents the bifurcation point (Ponce-Alvarez & Deco, 2024). Here, we used a slightly negative *a*_*j*_ value very close to the bifurcation point (*a*_*j*_ = −0.02), since it has been shown to represent an optimal working point for fitting whole-brain neuroimaging data (Deco et al., 2017b). Finally, the fMRI signals were modelled by the real part of the state variables, i.e. *x*_*j*_= *Real*(*z*_*j*_).

### Linearisation of the model

The proximity to criticality given by the bifurcation parameter allows the linearisation of the dynamics around the fixed point ***z*** = 0, which for *a* < 0 is stable (Ponce-Alvarez & Deco, 2024). This is essential because it allows us to find an analytical solution for the functional connectivity matrix **C**. The functional correlations between all pairs of brain regions can be estimated by employing a linear noise approximation (LNA). Consequently, the dynamical system of *N* nodes (**Equation 5**) can be expressed in a vector format as follows:

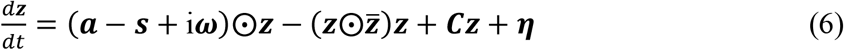

where ***z*** = [*z*_1_, …, *z*_*N*_]^*T*^, ***a*** = [*a*_1_, …, *a*_*N*_]^*T*^, ***ω*** = [*ω*_1_, …, *ω*_*N*_]^*T*^, ***η*** = [*η*_1_, …, *η*_*N*_]^*T*^, ***s*** = [*s*_1_, …, *s*_*N*_]^*T*^ is the vector containing the strength of each node, i.e., *s*_*i*_ = ∑_*j*_ *C*_*ij*_, [ ]^*T*^ represents the transpose and ⨀ is the Hadamard element-wise product, i.e., **u**⨀**v** = [u_1_v_1_, …, u_*N*_v_N_], and 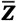 is the complex conjugate of ***z***.

Linearisation allows dropping high order terms in **Equation 6** and the dynamics of linear fluctuations can be modelled using a Langevin stochastic differential equation

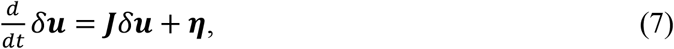

where fluctuations of ***x*** and ***y*** are described by a 2N-dimensional vector *δ****u*** = [*δ****x***, *δ****y***]^***T***^ = [*δx*_1_, …, *δx*_*N*_, *δy*_1_, …, *δy*_*N*_]^*T*^, ***η*** is a white noise 2N-dimensional vector, and ***J*** is the 2N × 2N Jacobian of the system evaluated at the fixed point (***x, y***) = (**0, 0**), given by

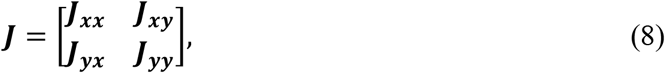

where ***J***_***xx***_, ***J***_***xy***_, ***J***_***yx***_, ***J***_***yy***_ are N × N matrices given by ***J***_***xx***_ = ***J***_***yy***_ = ***C*** + *diag*(***a*** − ***s***) and ***J***_***yx***_ = − ***J***_***xy***_= *diag*(***ω***), being *diag*(***v***) a diagonal matrix with a vector ***v*** on the diagonal (Deco et al., 2024; Ponce-Alvarez & Deco, 2024).

This equation helps to find the temporal evolution *δ****u***, from which network statistics can be extracted. For instance, the covariance of fluctuations 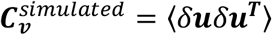 is here relevant, since it will allow to define simulated FC. From **Equation 7**, the differential equation governing ***C***_***v***_ can be deduced

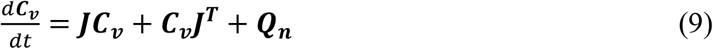

where ***Q***_*n*_ is the noise covariance matrix *Q*_*n*_ = ⟨***ηη***^***T***^⟩. If 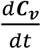 is set to zero, the stationary covariance can be obtained by solving:

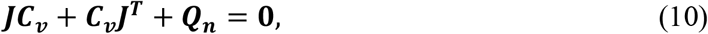

This is a Lyapunov equation that has a unique solution provided that ***J*** is asymptotically stable. It can be solved using the eigen-decomposition of the Jacobian matrix ***J*** (Deco et al., 2014; Ponce-Alvarez & Deco, 2024). Here, it was solved using MATLAB solver *lyap()*.

Note that the stationary covariance gives a measure of how fluctuations from different regions covary at stationarity, i.e., when the stable origin ***z*** = **0** is reached. The stationary covariance can be used to calculate the forward *τ* time-shifted covariance, defined as 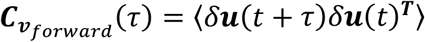, using:

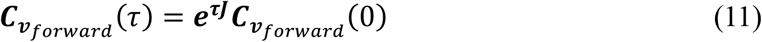

where 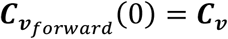 is the static covariance (i.e., with zero shift). For deeper mathematical insight, see Ref. (Ponce-Alvarez & Deco, 2024). Since we are only interested in the real part of network statistics, simulated covariance 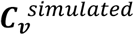 and simulated forward time-shifted covariance 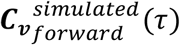 matrices will refer to the matrices arising from taking the first N rows and N columns of ***C***_***v***_ and 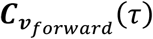, respectively. Analogously to empirical measures, simulated static functional connectivity ***FC***^*simulated*^and forward *τ* time-shifted functional connectivity 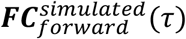 can be obtained with:

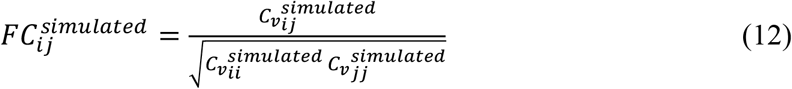

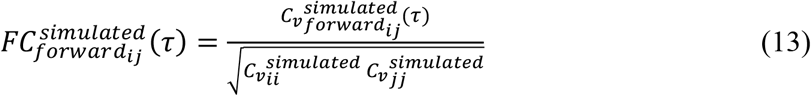

### Model fitting: Generative Effective Connectivity

Given a coupling connectivity matrix ***C***, the Hopf whole-brain model simulates the local dynamics of the parcellated brain regions, enabling the computation of both static and time-shifted simulated FC matrices through network statistic measures of real-part fluctuations. Here, patient-specific SC matrices were not available, so ***C*** matrices were initialised to a group SC template from the Human Connectome Project (Deco et al., 2021). More specifically, in order to ensure consistency with prior literature set of parameters, ***C*** was set as a scaled version of the template, ensuring a maximal ***C*** value of 0.2 (Deco et al., 2017b). Models were then fitted to the empirical data by optimising ***C*** following a pseudo-gradient procedure. At each iteration, ***C*** weights were refined according to the following updating rule:

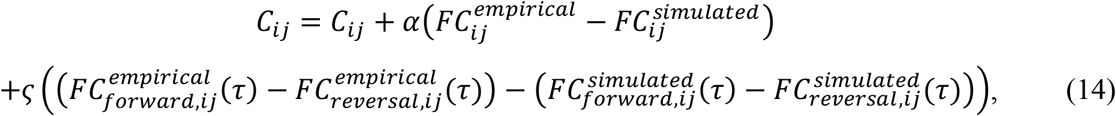

where 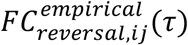 and 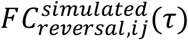 denote the empirical and simulated reversal *τ* time-shifted FC, analogously computed as the forward time-shifted counterparts but considering the time reversal of the empirical and simulated time series. The learning rates used were *α* = 0.0004 and *ς* = 0.0001 were used.

Note that maximal resemblance between 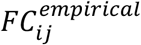 and 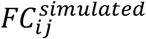 was sought, but also between 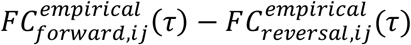 and 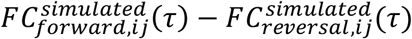, both matrices of temporal asymmetry which estimate the pairwise levels of temporal irreversibility between time series (Deco et al., 2022). Since temporal irreversibility is linked to the hierarchy of causal interactions in brain dynamics (Deco et al., 2022; Kringelbach et al., 2023), including it in model optimisation is essential to encode brain hierarchy in GEC matrices. Note also that fitting the time asymmetry matrices is equivalent to fitting only the forward time shifted correlations.

To check the quality of the fitting, every 100 iterations of the group and subject-level optimisation process, a total fitting error *ϵ*_*total*_ was calculated, given by:

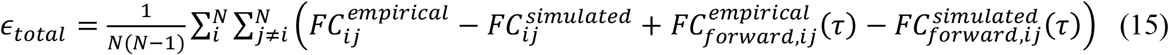

The optimisation loop was exited whenever the error varied less than 0.1% with regards to the previous error computed. The resulting optimised ***C*** matrix is called Generative Effective Connectivity (GEC) and contains the effective conductivity values for each existing pair of anatomical connections and not just the dMRI based density of fibres (Deco et al., 2024).

To accelerate the fitting process, ***C*** optimisation was divided in two stages. First, ***C*** was fitted to group average empirical FC matrices, obtaining group GECs for each of the defined conditions: psilocybin arm before treatment (PB), psilocybin arm after treatment (PA), escitalopram arm before treatment (EB) and escitalopram arm after treatment (EA). Second, group GECs were used as initial ***C*** matrices to fit patient-specific models, using patient-specific empirical data. The success of model fitting was tracking the total fitting error and computing the correlation between ***FC***^*empirical*^ and ***FC***^*simulated*^ as well as between 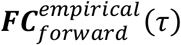 and 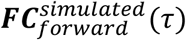. **Figure 1** shows a graphical representation of the pipeline for GEC generation via whole-brain modelling.

**Figure 1.**
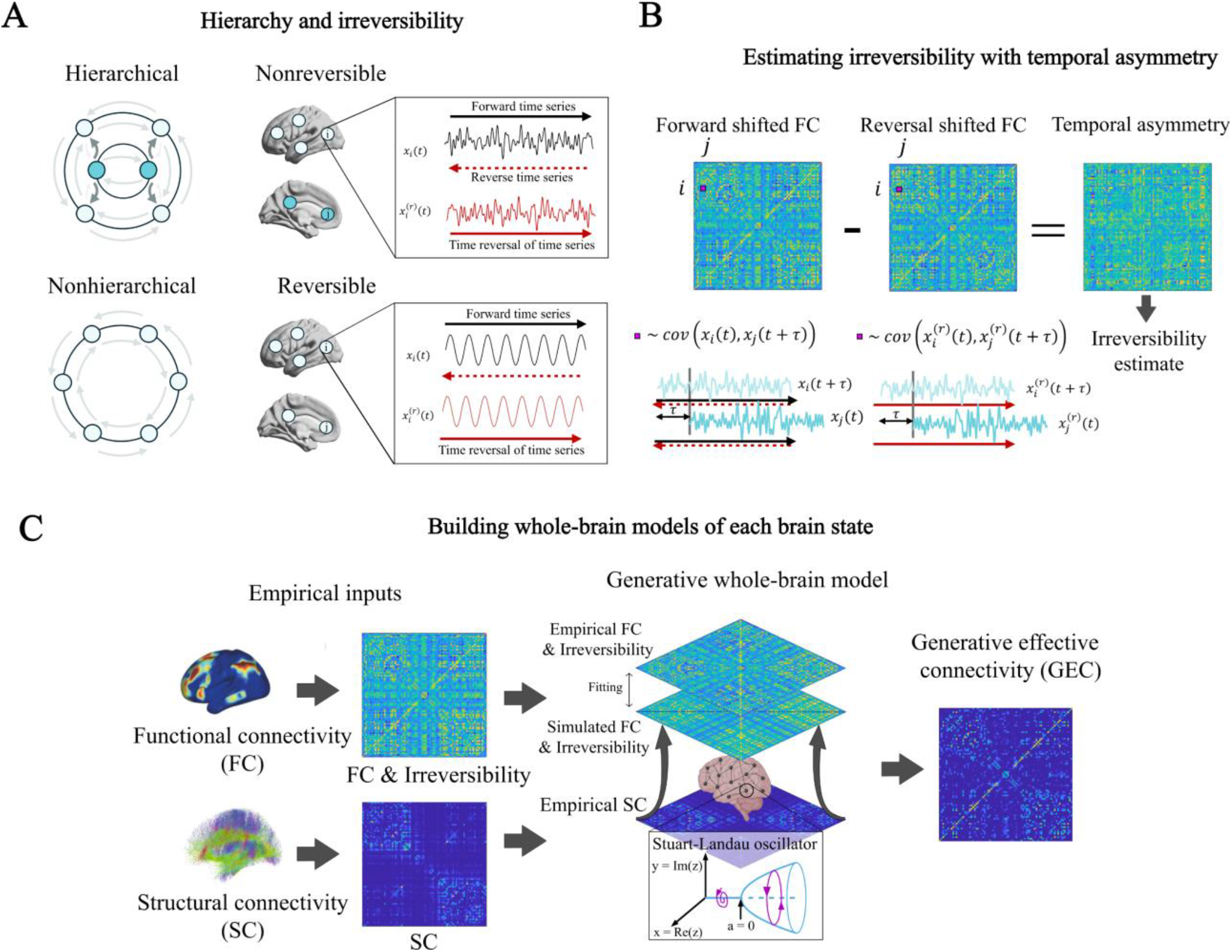
Thermodynamics of mind framework. The figure shows how the principles from thermodynamics can be used to infer the hierarchy of brain states. **A)** The level of hierarchy in a brain network is deeply connected to temporal irreversibility of the produced neural time series. In hierarchical networks, information flows asymetrically between its nodes, with intrinsically-driven nodes (turquoise) driving the (extrinsic) activity of lower regions (light blue). Thermodinamically, they break the detailed balance, establishing an arrow of time. In other words, the produced time series become non-reversible. Visually, we can assess the irreversibility of a signal by temporally reversing its time series and inspecting its plausibility. Nonhierarchical networks, instead, do not break the detail balance, making the generated signals reversible in time. **B)** The irreversibility of a network can be estimated by constructing a matrix of the temporal asymmetry between regional time series. For each interaction, temporal asymmetry can be defined as the difference between forward and reversal time-shifted correlations. **C)** Whole-brain modelling can be used to identify the causal, mechanistic generators of functional hierarchy. The model uses local dynamics (for example, Stuart-Landau oscillators), initially informed by the empirical structural connectivity to fit the empirical functional connectivity and irreversibility. The optimisation adjusts the weights of the anatomical connections, producing a generative effective connectivity (GEC) that estimates the generators of the functional hierarchy.

### Single-site perturbation protocol

Upon computing GEC networks for both sessions in each patient, we started perturbing the corresponding ―fitted― whole-brain models. Perturbed models were those arising from the optimisation process, with GEC informing the asymmetric connectivity between nodes. In-silico single-site perturbations consisted of increases in the white noise standard deviation *β* of a particular Stuart-Landau oscillator (see **Equation 5**). Such disturbance can be interpreted as pushing the oscillations to a more noisy-driven regime. We also refer to the noise level *β* in this context as the *perturbation* or *stimulation intensity* and we call *perturbation* or *stimulation target* the region or site corresponding to the perturbed oscillator. All models underwent a systematic single-site perturbation protocol, meaning that for every model, all nodes were individually perturbed with a repertoire of intensities ranging from 0.02 to 0.5 in steps of 0.01. Each perturbed whole-brain model generated a perturbed simulated FC, capturing the functional statistical dependencies between regions under perturbation. Two measures were constructed to assess the impact of each perturbation: susceptibility and Perturbation Effectivity for Recovery.

### Susceptibility to perturbation

Susceptibility (S) aims to inform on the brain’s *capacity to change* its functional patterns under a given perturbation. Consequently, we defined susceptibility as a dissimilarity measure between perturbed and unperturbed FCs, assessed in terms of decorrelation:

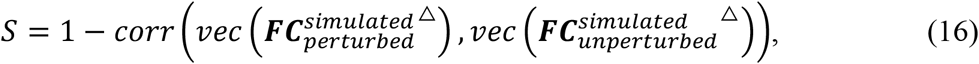

where *corr*( ) stands for Pearson’s correlation and *vec*(***A***^△^) denotes the vector formed by the upper triangular elements of the matrix **A**. Note that only the upper triangular elements are needed, given the symmetry of FC matrices and the trivial values of self-connections (ones) in any correlation-based FC matrix (Liégeois et al., 2020).

To compare the susceptibility present in psilocybin and escitalopram-induced brain states we studied, for each patient, the average susceptibility across all possible targets. Note that for such computation we fixed the stimulation intensity, given that its influence in susceptibility.

### Perturbation Effectivity for Recovery

Susceptibility, which captures functional malleability, has the limitation of not reporting whether the change exerted by perturbations is beneficial or not, here understood as enhancing a transition towards a healthier regime. The assessment of in-silico stimulations was therefore complemented with another measure, termed Perturbation Effectivity for Recovery (PER). This concept has been previously described in other computational studies (Vohryzek et al., 2023b), and aims to capture the effectivity of perturbations in driving the brain towards a healthy state. We constructed a healthy functional target by averaging the resting state FC of 100 healthy participants. As mentioned earlier, fMRI DK80-parcellated signals from 100 unrelated subjects part of the HCP Young Adult Project were accessed, and empirical FCs were computed as shown in **Equation 2**. In contrast with susceptibility, PER was defined as a correlation measure:

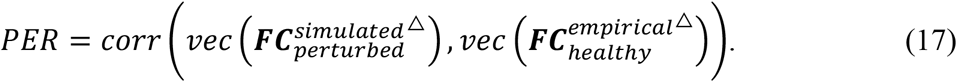

Furthermore, to strictly measure the effect that perturbations have in driving healthy transitions, we must consider the baseline similarity of unperturbed FCs to the healthy target. We call this baseline Baseline Similarity to Recovery (BSR) and define it by:

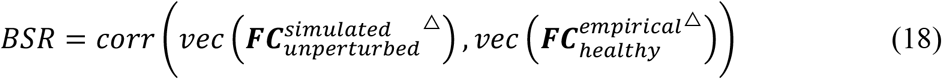

By removing the baseline from PER, we obtain the normalised Perturbation Effectivity for Recovery (nPER) measure:

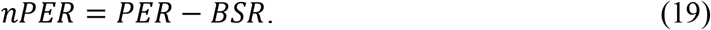

Perturbations were considered to drive a healthy transition whenever nPER was positive, i.e., whenever perturbations increased the similarity between the simulated FC and the healthy target. Since we want to elucidate the differences between psilocybin and escitalopram treatments in terms of how beneficial can perturbations become, we focused on identifying the healthiest transition achievable for each patient in each group―across all regions perturbed and all stimulation intensities applied―.Mathematically, this was done by determining the maximum PER in each patient (i.e.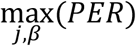). The corresponding stimulation site and stimulation intensity were collectively referred nto as the patient’s *optimal perturbation*. The general framework of the perturbation protocol and how susceptibility and PER can be used to assess brain states under perturbation is illustrated in **Figure 2**.

**Figure 2.**
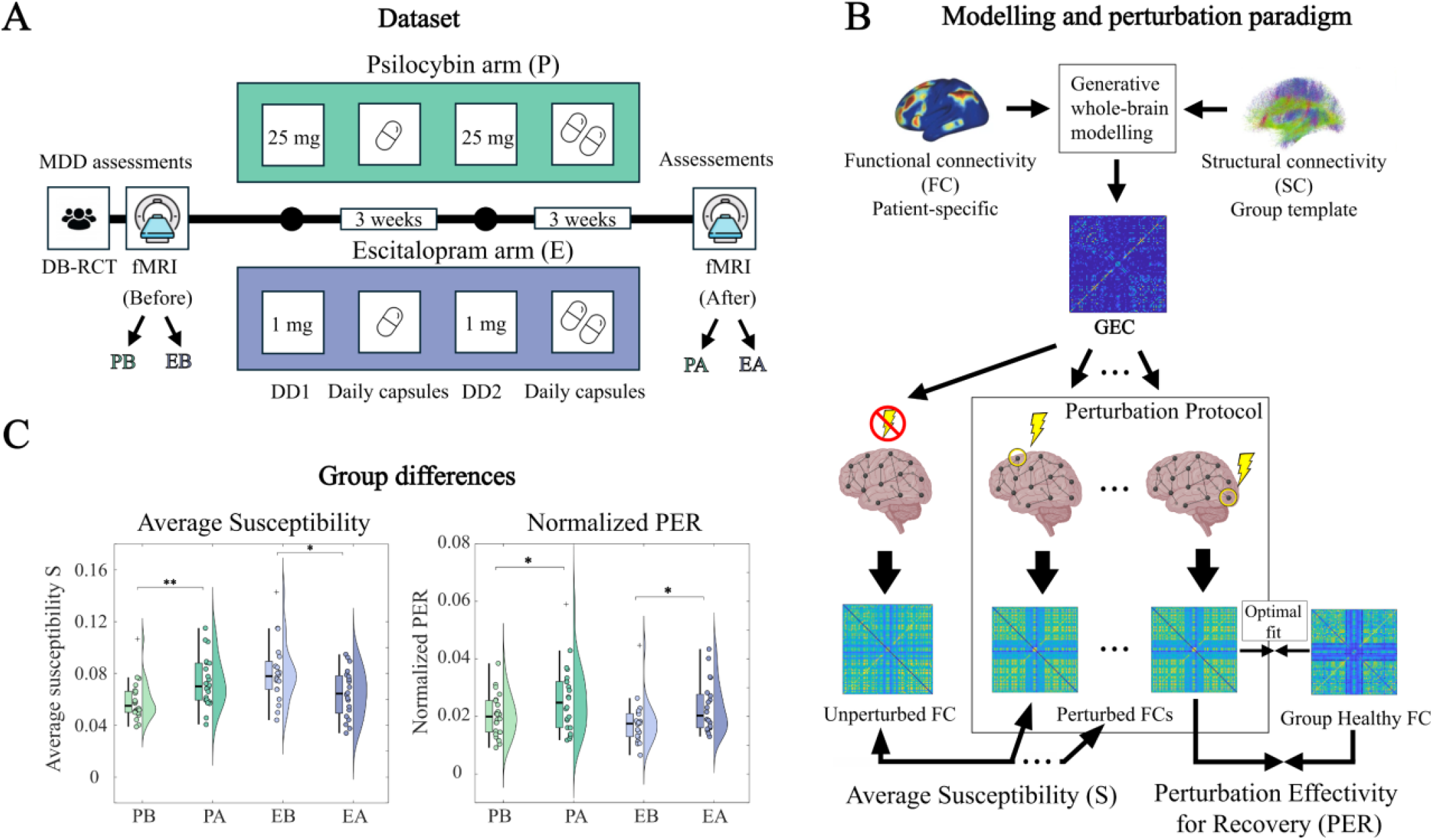
Perturbing whole-brain models of depression under different pharmacological interventions. **A)** Neuroimaging resting state data was acquired before and after a double-blind phase II randomised controlled trial comparing psilocybin therapy with escitalopram. **B)** Whole-brain models were built and perturbed for each treatment arm, before and after pharmacological intervention. For each patient, a whole-brain model was first fitted to the empirical FC data via the optimisation and generation of a Generative Effective Connectivity (GEC) matrix capturing the asymmetric effective conductivity between regions. Then, patient whole-brain models informed by GECs were systematically perturbed, leading to a perturbed FC. We defined susceptibility as a dissimilarity measure between unperturbed and perturbed simulated FC, and Perturbation Effectivity for Recovery as a similarity measure between perturbed simulated FC and a healthy FC target. **C)** Statistical tests of susceptibility and PER between treatment arms before and after treatment elucidate group differences.

### Multi-site perturbation protocol

While asking the question of how pharmacological treatments boost the therapeutical benefits of neurostimulation for depression, it is reasonable to explore the simultaneous stimulation of multiple regions rather than constraining our in-silico protocols to single-site approaches. After all, depressive states have been linked to aberrant activity in brain networks like the default-mode network (Menon, 2011), suggesting that multi-site protocols might outperform local stimulations at single regions. To explore this, we employed a multi-site stimulation protocol. Given that the number of possible multi-site configurations scales exponentially with the number of regions considered, we employed a greedy perturbation algorithm, which constructed optimal configurations by iteratively adding the single-site perturbations that jointly maximise PER.

The greedy level was set to 20 areas, and the algorithm run using PER maximisation. For each greedy level, all eligible areas ―all of them for the first level― were perturbed with a stimulation intensity repertoire ranging from 0.01 to 0.1 in steps of 0.01. The pair of area and noise level showing the highest PER value was selected, with that area being removed from the eligible set. Note that noise levels above 0.1 were here not considered to reduce computation time, after realising that the highest PER values occur for rather small perturbations (**Figure S5**).

### Statistical analysis

For a given measure of interest, statistical tests were performed to assess statistical significance between groups. Essentially, such tests involved assessing the effects of treatment (before vs after), as well as comparing differences between treatment arms (psilocybin vs escitalopram). Due to the relatively small sample size (22 patients for psilocybin and 20 for escitalopram), non-parametric Wilcoxon tests were conducted, not assuming Gaussian distributions in the studied distributions. Paired Wilcoxon tests were used to study treatment effects, while unpaired Wilcoxon tests were employed to assess differences between arms. For both cases, Wilcoxon tests were implemented using 5000 permutations to approximate the null hypothesis distribution from the data. Statistical significance was declared for p-values below 0.05 (p < 0.05). In the figures, three different levels of significance are denoted using the following asterisk code: *** for p < 0.001, ** for p < 0.01, and * for p < 0.05. Moreover, we refer with PB, PA, EB and EA to the conditions *psilocybin arm before treatment, psilocybin arm after treatment, escitalopram arm before treatment*, and *escitalopram arm after treatment*, respectively.

## Results

In this work, we investigated the effects of perturbations in the depressed brain following psilocybin and escitalopram treatments. More specifically, we analysed the susceptibility of functional brain states to perturbations and their drivability to optimal states following both pharmacological interventions. The dataset analysed originates from a double-blind phase II randomised controlled trial comparing psilocybin and escitalopram, where resting-state fMRI scans were taken before and after pharmacological intervention, as shown in **Figure 2A** (see details of the trial in *Materials and Methods*).

### Generative whole-brain modelling and brain hierarchy

For each patient scan, whole-brain modelling, constrained by the structural connectivity, enabled the estimation of a generative effective connectivity (GEC), capturing the resting-state brain dynamics. Following the DK80 parcellation, whole-brain models comprised 80 Stuart-Landau oscillators, and they were initially informed by a structural connectivity (SC) template, capturing the anatomical connections between regions. These connections were iteratively updated to make the simulated activity fit the patient-specific empirical data, which included static functional connectivity (FC) and estimates of pairwise temporal irreversibility. The latter was motivated by the Thermodynamics of Mind framework, which links pairwise temporal irreversibility between neural time series to functional brain hierarchy (Kringelbach et al., 2024). Here, temporal irreversibility was estimated using temporal asymmetry, defined by the difference between forward and reversal time-shifted correlations and in line with the INSIDEOUT framework (Deco et al., 2022). By including estimates of pairwise irreversibility in the fitting process, the produced GEC captures the hierarchy of the causal influences between regions. **Figure 1A** shows the relationship between hierarchy and irreversibility. Namely, in hierarchical networks, the reversed time series of neuronal activity becomes highly implausible, whereas in networks with a flat hierarchy, neuronal activity is reversible. **Figure 1B** illustrates how temporal irreversibility can be estimated using temporal asymmetry. Last, the process of employing generative whole-brain models to capture the hierarchy of specific brain states is shown in **Figure 1C**.

### Whole-brain modelling under perturbation

Following the concept of Dynamic Sensitivity Analysis, optimised whole-brain models informed by patient-specific GECs were then systematically perturbed by increasing the amount of noise in the local oscillations of perturbed nodes. The aim was, in the context of depression, to figure out under which brain states can the brain benefit more from external perturbations, treating the resulting perturbed functional connectivity as the output of interest. We first ran single-site perturbations in all nodes with a range of stimulation intensities, and later explored the power of multi-site perturbation. We constructed two measures to assesses the impact of perturbations on functional connectivity: susceptibility (S) and Perturbation Effectivity for Recovery (PER). The former measured the dissimilarity of the perturbed FC with respect to the unperturbed FC, reporting hence the capacity of the brain’s activity to change under perturbation, and the latter measures the proximity of the perturbed FC with respect to a healthy functional target, extracted from the average FC of 100 healthy unrelated participants (Vohryzek et al., 2020). **Figure 2B** illustrates the perturbation protocol as well as how susceptibility and PER measures were defined. By performing statistical analyses, we were able to quantify the effects of psilocybin and escitalopram treatments in these measures (**Figure 2C**).

### Susceptibility to perturbations

Susceptibility (S) was computed for all single-site perturbations. To extract statistics across patients, we computed the average susceptibility across all perturbed targets (fixing the stimulation intensity) for the different groups (**Figure 3A**). Results show that for a stimulation intensity *β* = 0.04, psilocybin significantly increases susceptibility (p<0.01, paired Wilcoxon test with 5,000 permutations) whereas escitalopram significantly decreases it (p<0.05, paired Wilcoxon test with 5,000 permutations). The significance of this trend was analysed across stimulation intensities, showing that the psilocybin trend is stronger than escitalopram’s one and that significance is guaranteed for intensities between 0.02 and 0.05 (**Figure S3**). **Figure 3B** shows the number of areas increasing susceptibility following psilocybin and escitalopram treatments. In line with prior results, psilocybin proves to increase susceptibility in significantly more areas than escitalopram (p<0.001, unpaired Wilcoxon test with 5,000 permutations), where perturbations rather tend to decrease it. Finally, the prevalence of susceptibility increases across patients is rendered region-wise for each perturbed area (**Figure 3C**).

**Figure 3.**
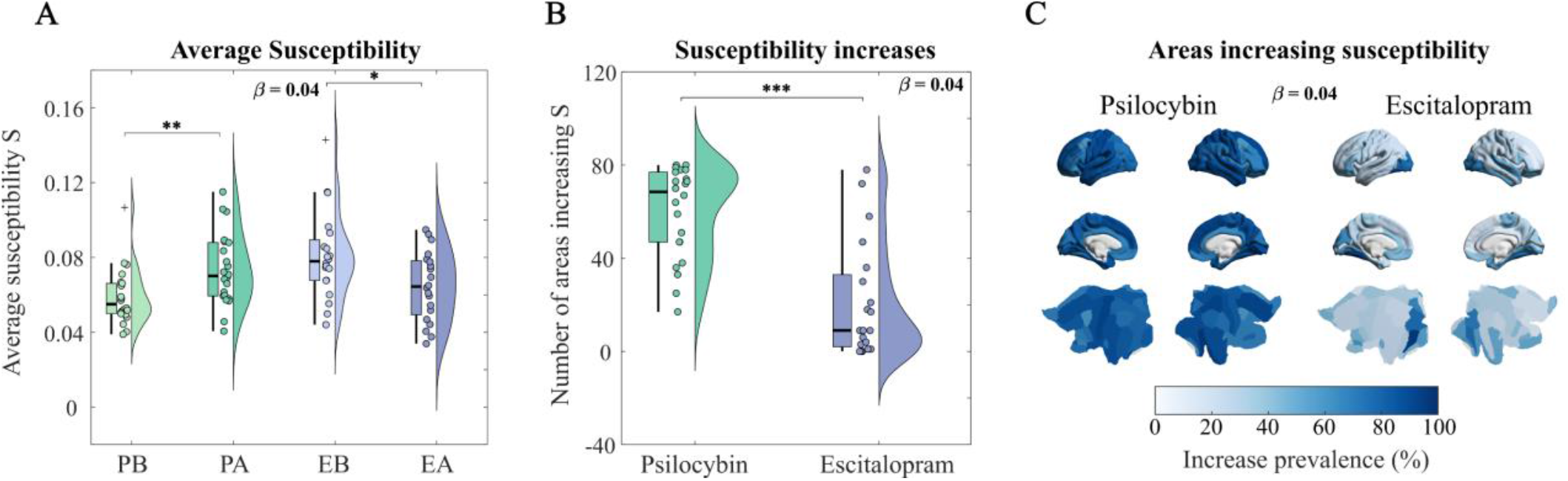
Opposite pharmacological reconfigurations in susceptibility. **A)** For a fixed stimulation intensity β = 0.04, the average susceptibility across perturbation sites is shown for each group. Psilocybin significantly increases average susceptibility (^**^p<0.01, paired Wilcoxon test with 5,000 permutations) whereas escitalopram significantly decreases it (^*^p<0.05, paired Wilcoxon test with 5,000 permutations). **B)** The number of areas increasing susceptibility is plotted for each treatment arm. Psilocybin significantly differs from escitalopram treatment (^***^p<0.001, unpaired Wilcoxon test with 5,000 permutations), with a tendency to increase susceptibility in more areas. **C)** The renderings show the areas that increase susceptibility more consistently across patients.

### Perturbation Effectivity for Recovery

As with susceptibility, Perturbation Effectivity for Recovery (PER) was computed for every single-site perturbation. Here, however, the interest lied in identifying the perturbation (intensity and location) with highest PER, i.e. driving the simulated FC the closest to the healthy target. **Figure 4A** shows the distributions of PER in opposition to the Baseline Similarity to Recovery (BSR) for each group and session. For all groups, PER values are significantly higher than BSR (p<0.001, paired Wilcoxon tests with 5,000 permutations), proving that if properly tailored, perturbations significantly improve the similarity to the healthy state. **Figure 3B** shows the distributions of normalised PER (nPER), obtained by taking out BSR from PER. Psilocybin and escitalopram treated brains show significantly higher nPER values for their best perturbations (p<0.05, paired Wilcoxon tests with 5,000 permutations), showing that pharmacological treatment enhances the capacity to benefit from perturbation. Finally, **Figure 3C** shows nPER distributions for multi-site perturbations in opposition to the single-site approach. Significantly higher nPER values are achieved by multi-site perturbations (p<0.001, paired Wilcoxon tests with 5,000 permutations), proving that multi-site perturbations can be more beneficial than best single-site solutions if the set of stimulation targets and intensities is properly selected.

**Figure 4.**
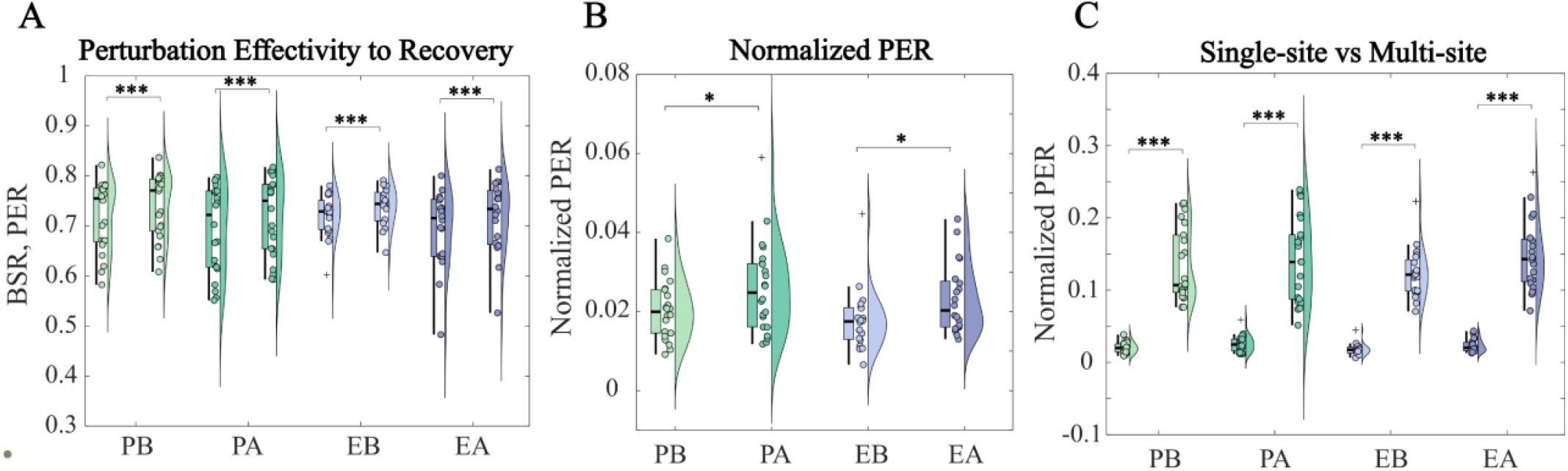
Healthier transitions unlocked by psilocybin and escitalopram. **A)** The figure shows, for each group and condition, the Baseline Similarity to Recovery (left) versus the highest Perturbation Effectivity for Recovery (left) achieved in each patient. These distributions remain significantly different for all groups and conditions (^***^p<0.001, paired Wilcoxon tests with 5,000 permutations). H**B)** The normalised Perturbation Effectivity for Recovery is plotted for each group and condition. Both psilocybin and escitalopram significantly increase this quantity (^*^p<0.05, paired Wilcoxon tests with 5,000 permutations). **C)** The highest nPER of single-site perturbations is plotted (left) against the nPER of the 20-level-greedy-best multi-site perturbation. Multi-site perturbation significantly reaches higher nPER values (^***^p<0.001, paired Wilcoxon tests with 5,000 permutations).

### Most optimal stimulation targets

We finally examined the best stimulation targets following pharmacological intervention, according to best single-site and greedy-best multi-site stimulations (**Figure 5**). **Figure 5A** shows the prevalence across patients of best single-site stimulation targets, coloured on coronal and axial slices of the brain. The exact prevalence can be observed in **Figure 5B**. Overall, we found that the right amygdala is the best stimulation target, present in 50% of best single-site perturbations following both treatments. The nucleus accumbens covered the other 50%, split between right and left areas. **Figure 5C** shows the prevalence across patients of 20-level greedy multi-site stimulation targets, coloured on coronal and axial slices of the brain. **Figure 5D** shows the same information in a bar plot, but only considering areas with a prevalence higher than 30%. The most frequently recruited areas are again the right and left nucleus accumbens as well as the right amygdala. Moreover, the right medial orbitofrontal cortex is the most frequently recruited cortical target.

**Figure 5.**
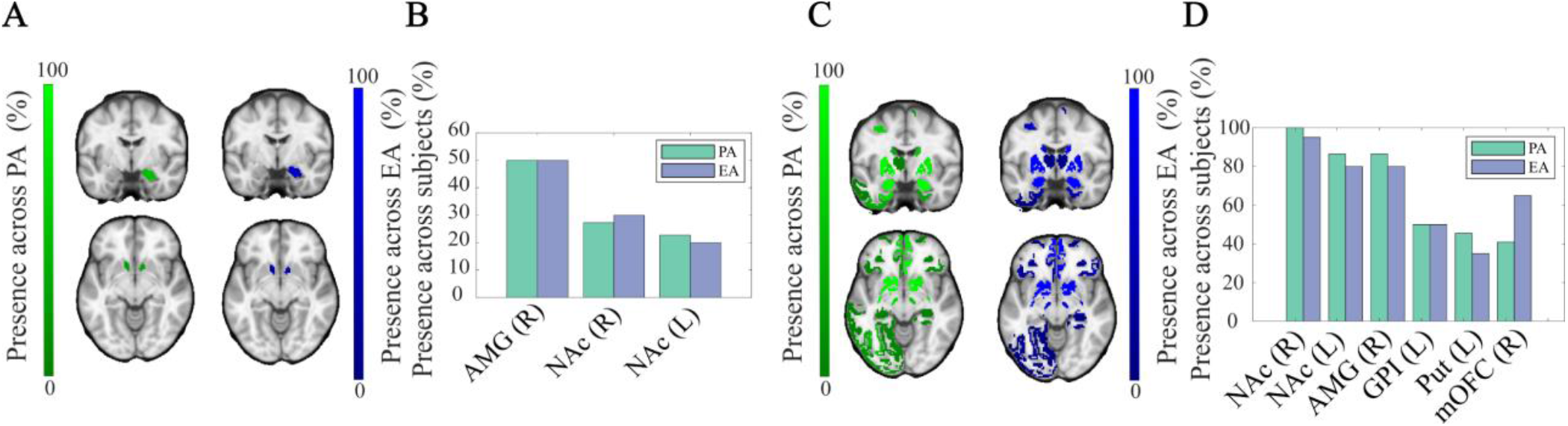
The nucleus accumbens and the right amygdala as optimal targets in single- and multi-site stimulations. **A)** The presence of stimulation targets in optimal single-site perturbations is shown in coronal (top) and axial (bottom) slices of the brain following psilocybin (left) and escitalopram treatments. **B)** The bar plot displays the same information than A). **C)** The presence of stimulation targets in 20-level-greedy multi-site perturbations is shown in coronal (top) and axial (bottom) slices of the brain following psilocybin (left) and escitalopram treatments. **D)** The bar plot shows the same information than C) for those areas showing a presence higher than 30%.

## Discussion

In this work, our aim was to study in-silico the susceptibility of brain states to perturbations and their drivability to optimal states following psilocybin and escitalopram treatments. Our hypothesis was that psilocybin should enhance susceptibility and improve the drivability to healthier brain states, whereas escitalopram should on average constrain functional malleability but still enhance healthy transitions for specific stimulations. We addressed our research question by first fitting whole-brain models to the patient’s empirical data, and then systematically perturbing the fitted models by increasing the noise in the modelled local dynamics. Single-site and multi-site stimulation protocols were explored, and we assessed their impact on static functional connectivity via two distinct measures: susceptibility, which quantifies the ability of perturbations to change the FC pattern, and Perturbation Effectivity for Recovery (PER), which measures the proximity of perturbed FC to a healthy target (where a normalised PER was also defined strictly measuring the impact of perturbations in driving healthy transitions).

Overall, our findings suggest that psilocybin enhances the susceptibility of brain states to perturbations and improves their drivability to optimal states, whereas escitalopram generally constrains functional malleability but still facilitates healthy transitions under specific stimulations. We also demonstrate that the most optimal stimulation targets to achieve healthy transitions occur in subcortical areas like the amygdala and the nucleus accumbens, and we show that multi-site stimulation protocols can significantly outperform single-site strategies. Crucially, our analysis involved systematically perturbing whole-brain models that were fitted not only to patients’ static functional connectivity but also to an estimate of their functional hierarchy. Given that psilocybin and escitalopram have been associated with different brain hierarchical reconfigurations, the present methodology constitutes a promising avenue for the in-silico assessment of brain state transitions following pharmacological treatment for depression, with potential applications in other neuropsychiatric disorders.

### Opposite overall reconfiguration in susceptibility by psilocybin and escitalopram

The analysis of susceptibility in single-site perturbations revealed an opposite pharmacological trend: psilocybin increases the average susceptibility to perturbations (for a fixed stimulation intensity), whereas escitalopram diminishes it (**Figure 3A**). For this analysis, we crucially compared single-site stimulations under a same fixed stimulation intensity, since we wanted to study the exclusive dependency of susceptibility on pharmacological treatment (the dependency of susceptibility on stimulation intensity can be observed in **Figure S4**). In line with the first result, we also found that the number of areas where perturbation’s susceptibility increases with treatment is larger in the psilocybin arm than in the escitalopram arm, where most single-site perturbations diminish susceptibility (**Figure 3B-C**). This further confirms the opposite reconfiguration exerted by psilocybin and escitalopram in susceptibility to perturbation.

Such an opposite effect aligns with the known differential effects on pharmacology of the two drugs. On the one hand, psilocybin is known to act mainly through the serotonin 2A receptor (5-HT_2A_R) (Kringelbach et al., 2020; Nichols, 2016; Vargas et al., 2023), initiating multi-level plasticity events (Carhart-Harris, 2019). On the other hand, escitalopram is a selective serotonin reuptake inhibitor (SSRI). These drugs constitute the first-line pharmacological treatment for depression and are thought to act on the serotonin transporter, where psilocybin has no appreciable affinity (Chu & Roopma, 2023).

By how susceptibility was defined, we interpret its increase following psilocybin treatment as an opening of a plasticity window (Carhart-Harris & Nutt, 2017; Ruffini et al., 2024b). Conversely, the diminishment observed in the escitalopram arm suggests that escitalopram constrains functional malleability, limiting the brain’s capacity to change under perturbations. As motivated in the introduction, these results align with a recent computational study where differential reconfigurations on the brain’s overall hierarchy were associated to psilocybin and escitalopram (Deco et al., 2024). Specifically, the authors used novel topological measures adapted to directed networks to show that psilocybin flattens the brain hierarchy, whereas escitalopram increases it. Bringing it together with the present study, we demonstrate that the flattening of functional hierarchy can be related to increases in susceptibility to model perturbations, and that conversely, enhanced hierarchies difficult the ability of brain functional activity to change, on average, to local perturbations. This further supports the REBUS model, where psychedelics would open a therapeutic window of plasticity by flattening the brain’s hierarchy (Carhart-Harris & Friston, 2019).

### The optimal transitions are found after both interventions

Systematic single-site perturbations were further assessed in terms of their ability to recover healthy functional connectivity patterns, quantified through PER values. We first showed that perturbations, if properly tailored in stimulation site and intensity, lead to significantly higher PER values compared to the unperturbed Baseline Similarity to Recovery (BSR) values (**Figure 4A**). This was consistent across both treatment arms, before and after pharmacological intervention. This suggests that different pharmacological treatment interventions can meaningfully benefit from additional in-silico stimulations, highlighting the relevance of combined therapies where pharmacological intervention is accompanied by other treatments, like transcranial direct current stimulation (tDCS) or other neurostimulation solutions (Ruffini et al., 2024a; Ruffini et al., 2024b).

We then computed the normalised PER (nPER) values for each best perturbation, to strictly measure the influence of perturbations in driving healthier functional states (**Figure 4B**). Results demonstrate that after pharmacological intervention, nPER values significantly increase. This means that pharmacological intervention, whether psilocybin or escitalopram, enhances the therapeutic effectivity of optimal perturbations, suggesting a synergistic effect. In the case of escitalopram, this can seem counter-intuitive, especially after showing that it constrains, on average across sites, the ability of perturbations to impact FC. However, it is crucial to notice that even if susceptibility is generally decreased, some areas still experience susceptibility increases (**Figure 2B-C**). Since we are looking for the most optimal perturbation in each patient, the stimulation targets can and do correspond to such a minority of areas. Put together, escitalopram manages to reconfigure the brain in such a way that carefully selected perturbations in key regions still lead to meaningful healthy transitions, as meaningful as those found following psilocybin treatment. This aligns with the fact that both drugs manage to reduce depressive symptoms similarly (Carhart-Harris et al., 2021; Deco et al., 2024). Moreover, it is relevant to notice that the suitability of SSRIs for neurostimulation combined therapy has been largely studied, with some studies having found that the administration of serotonergic drugs like citalopram enhance the neuroplastic after-effects of anodal tDCS (Nitsche et al., 2009; Ruffini et al., 2024a). To this respect, our results suggest that escitalopram can enhance neurostimulation to drive functional patterns to healthier states, but since the functional malleability is constrained to the average perturbation, effective stimulations must target specific key regions. Last, the synergistic effect of psilocybin on unlocking healthier functional transitions may relate to the general increase in susceptibility. However, it is key to acknowledge that increased malleability can also imply increased vulnerability towards reaching unhealthy states. We here hypothesise that personalised stimulation protocols or supportive psychotherapy may represent the needed set of perturbations to effectively exploit the enhanced functional malleability, stimuli that might be too complex, especially in the case of psychotherapy, to be captured in this computational essay. Along with this idea, it might be that psilocybin is differentially beneficial, compared to escitalopram, in a longer-term after treatment completion, given that its neuroplastic-associated events can last for weeks (Calder & Hasler, 2023; Pasquini et al., 2024).

Given that depression alters brain function at the level of networks, i.e. by hyperactivating the default-mode network (DMN) (Menon, 2011), we further compared the therapeutic potential of multi-site perturbations with that of the single-site approach. Since the number of possible multi-site configurations scales exponentially with the number of considered nodes, we simplified the protocol by leveraging a 20-area greedy multi-site approach based on PER maximisation, after verifying that PER values saturate once approximately 20 areas are considered (**Figure S6**). As expected, we found that multi-site stimulations significantly reach higher nPER values (**Figure 4C**), indicating that healthier FC patterns can be restored. This result aligns with the body of research supporting bilateral for transcranial magnetic stimulation (TMS) (Mutz et al., 2019; Trevizol et al., 2019) as well as multichannel tDCS for the treatment of depression (Ruffini et al., 2024c), as well as the evidence showing that MDD disrupts large-scale brain networks (Menon, 2011). It is important to note, though, that our results concern also subcortical areas, which might be more challenging to access with non-invasive brain stimulation (NIBS) techniques like TMS or tDCS.

### The most optimal stimulation targets

Finally, potentially more relevant for the clinical setting, we analysed the targets where optimal perturbations occur following pharmacological intervention (i.e., where higher PER values are reached) (**Figure 5**). We found that following psilocybin and escitalopram treatments, these are either the right amygdala or the right and left nucleus accumbens (**Figure 5A-B**). The nucleus accumbens is known to be involved in the reward circuitry (Klawonn & Malenka, 2018) and to play a crucial role in MDD patients with anhedonia (Hu et al., 2023). Our results align with existing literature and support the recent interest in targeting this area through deep brain stimulation (Bewernick et al., 2010; Karrouri et al., 2021). Major depression has also been linked to alterations in the amygdala, which is associated with emotion regulation, among others (Perlman et al., 2019; Phillips et al., 2015). Our study suggests that stimulation in this area could notably improve depressive symptoms, a strategy that has been recently implemented using real-time fMRI neurofeedback (Young et al., 2017). The recruitment of only the right amygdala suggests lateralisation within the amygdala. In respect to this, the right amygdala is more commonly associated with automatic emotional processing, while the left amygdala is more linked to controlled mood regulation (Dyck et al., 2011; Murphy et al., 2020).

The targets recruited in 20-level greedy multi-site perturbations align with single-site perturbation results (**Figure 5C-D**), with the right amygdala, and the nucleus accumbens (left and right) showing the highest presence across patient configurations (following both psilocybin and escitalopram treatments). Other recruited areas are the left globus pallidus internus and, the left putamen, and the right medial orbitofrontal cortex (mOFC), which shows up as the best cortical target. This is especially relevant given the advantages of non-invasive brain stimulation (NIBS). Coherently with our results, mOFC is related to reward and mOFC reduced functional connectivity has been previously related to depression (Rolls et al., 2020).

Overall, this analysis shows that the most optimal stimulation targets to favour healthy transitions following psilocybin and escitalopram treatments are similar, and that best stimulation targets can vary across patients, strengthening the importance of personalised neurostimulation solutions, a concept already underscored for MDD given its heterogeneity (Gogulski et al., 2023).

### Limitations and future directions

The main limitation in this study is the scale of the data. FMRI data from only 42 patients was available (22 in the psilocybin arm and 20 taking escitalopram), a number that should be increased in further studies for increased validation of the results. Building and perturbing whole-brain models of larger datasets not only would better capture the heterogeneity associated to MDD (Malhi & Mann, 2018), but could also allow studying, for example, the ability of perturbation measures like PER to predict the depression severity after treatment.

With respect to future research directions, other perturbation protocols could be explored, for instance, by perturbing the bifurcation parameter in **Equation 5** instead of the noise level or by exploring more extensively the landscape of multi-site stimulations. Moreover, susceptibility and PER measures could be constructed in terms of FC dynamics and not just with a static average across time. For example, measures like metastability or Probable Metastable Substrates (PMS) could be used (Vohryzek et al., 2024a; Vohryzek et al., 2023b). This could provide meaningful complementary insights, especially given that psychedelics have been shown to broaden the repertoire of connectivity states across time (Atasoy et al., 2018; Cruzat et al., 2022; Tagliazucchi et al., 2014; Varley et al., 2020; Vohryzek et al., 2024b). Finally, this computational study could be complemented by examining fMRI signals some months after treatment, and not immediately upon treatment completion. Eventually, such lapse of time should allow the eventual benefits of longer-term plasticity events to show up in the different treated groups (Pasquini et al., 2024).

## Conclusion

To recapitulate, we show that psilocybin and escitalopram differentially reconfigure susceptibility to perturbations, with psilocybin increasing susceptibility on average across regions, and escitalopram decreasing it. This is consistent with the promise of psychedelics to open a window of plasticity from which supportive therapy could benefit and can be related to a flattening of the hierarchy of causal brain influences, as initially proposed by the REBUS model. We also show that healthier transitions can be promoted following the administration of both drugs. This suggests that neurostimulation can be equally useful following escitalopram treatment, but we hypothesise that the general increase of susceptibility with psilocybin may constitute a plastic advantage in a longer-term after the treatment is finished. Finally, we present the right amygdala and the nucleus accumbens as potential targets for stimulation, and we show that multi-stimulation significantly outperforms single-site approaches in driving healthy recovery of FC. Overall, the present work shows how Dynamic Sensitivity Analysis of whole-brain models of brain hierarchy can contribute to the in-silico personalisation of effective neurostimulation protocols for MDD, following or not pharmacological intervention.

## Acknowledgements

Jakub Vohryzek is supported by EU H2020 FET Proactive project Neurotwin grant agreement no. 101017716, Yonatan Sanz-Perl is supported by ‘ERDF A way of making Europe’, ERDF, EU, Project NEurological MEchanismS of Injury, and Sleep-like cellular dynamics (NEMESIS; ref. 101071900) funded by the EU ERC Synergy Horizon Europe, Morten L. Kringelbach is supported by the European Research Council Consolidator Grant: CAREGIVING (615539), Pettit Foundation, Carlsberg Foundation and Center for Music in the Brain, funded by the Danish National Research Foundation (DNRF117). Gustavo Deco is supported by the Spanish Research Project PSI2016-75688-P (Agencia Estatal de Investigación/Fondo Europeo de Desarrollo Regional, European Union); by the European Union’s Horizon 2020 Research and Innovation Programme under Grant Agreements 720270 (Human Brain Project [HBP] SGA1) and 785907 (HBP SGA2); and by the Catalan Agency for Management of University and Research Grants Programme 2017 SGR 1545. The neuroimaging analysis and whole-brain modeling is based on clinical research carried out at the National Institute for Health Research/Wellcome Trust Imperial Clinical Research Facility. The open-label trial was funded by a Medical Research Council clinical development scheme grant (MR/J00460X/1). The double-blind randomised controlled trial was funded by a private donation from the Alexander Mosley Charitable Trust, supplemented by Founders of Imperial College London’s Centre for Psychedelic Research. The funders had no role in study design, data collection and analysis, decision to publish or preparation of the paper.

## Author contribution

M.S-G., J.V., and G.D. designed the research; M.S-G., J.V., M.L.K., and R.C-H. performed the research; R.C-H. funded and oversaw the experimental research; M.S-G., J.V., Y.S-P., and G.D. contributed new analytic tools; M.S-G. analysed the data; and M.S-G., J.V., and G.D. wrote the paper.

