## Supplementary material for "Perturbing whole-brain models of brain hierarchy: an application for depression following pharmacological treatment": Perturbing whole-brain models of brain hierarchy - an application for depression following pharmacological treatment_SI_biorxiv.pdf

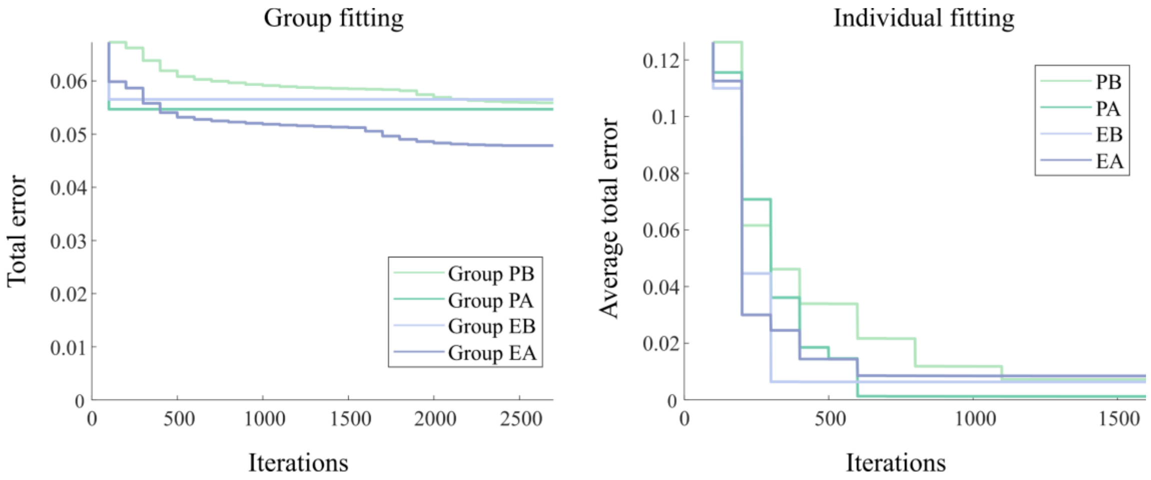


***Figure S1. Fitting total errors.*** *The temporal evolution of total errors for group (left) and individual fitting (right) is shown. For individual fitting, the average error across participants of the same group is shown. The error was arbitrarily initialized to 10000 and updated each 100 iterations until convergence (defined when the relative difference of the new error with respect to the previous error was less than 0.1%).*

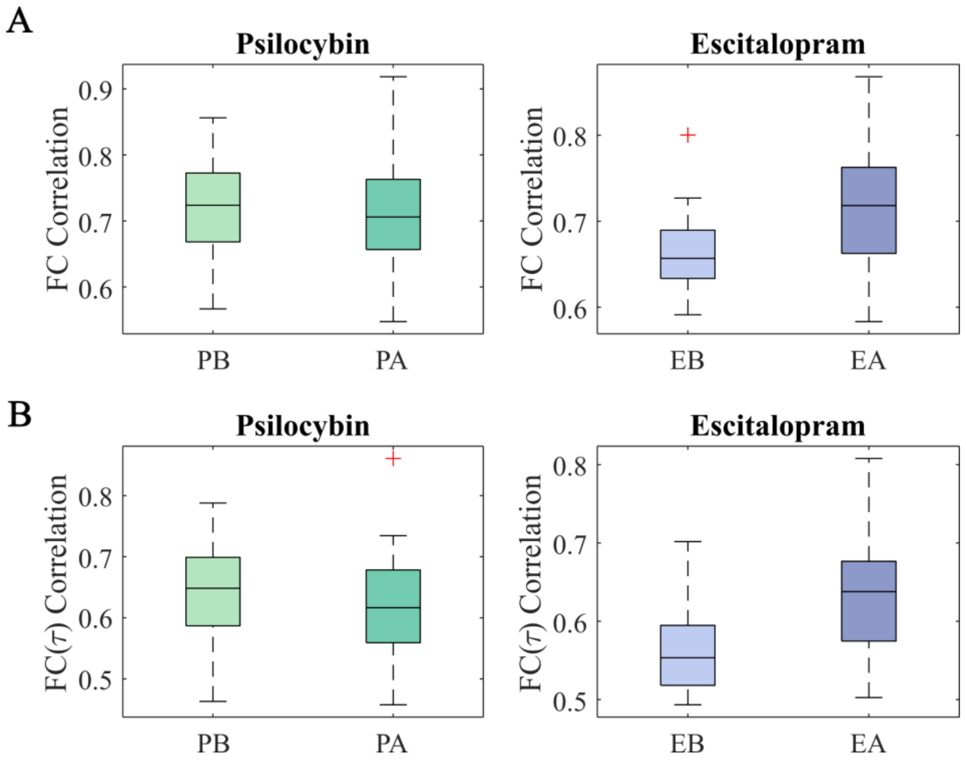


***Figure S2. Correlations between simulated and empirical data. A)*** *Correlations between simulated and empirical FC matrices are plotted, showing medians above 0.65 for both psilocybin (left) and escitalopram (right) groups before and after treatment (PB, PA, EB, EA).* ***B)*** *Analogous fitting correlations are shown using FS, exhibiting lower but still moderate correlation values for all groups.*

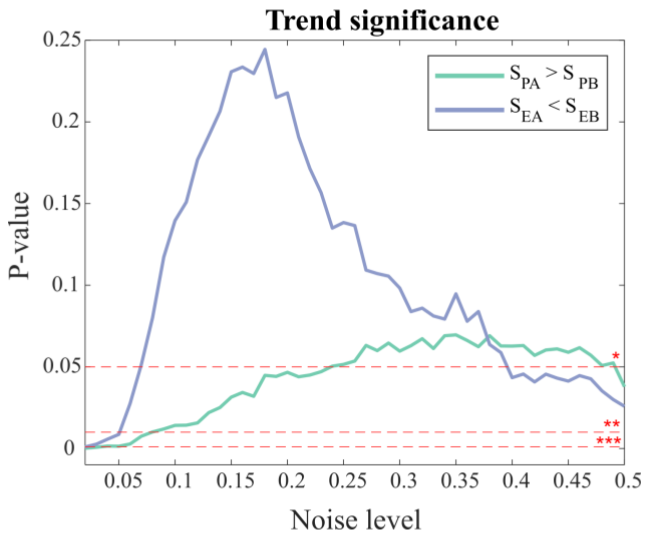


***Figure S3. Opposite pharmacological reconfigurations in susceptibility: Trend significance.*** *The plot shows the p-values corresponding to the probability that the differential effects observed following psilocybin and escitalopram treatment in susceptibility are found by randomly permuted data (paired Wilcoxon tests with 5000 permutations).*

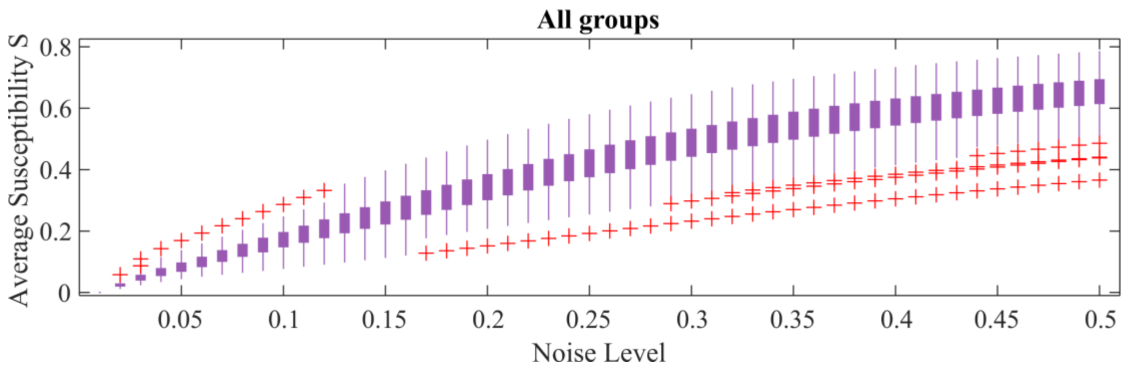


***Figure S4. Average susceptibility across noise levels.*** *The patient distribution of the average perturbation susceptibility across areas is plotted as a function of the noise level. The distributions represent all subjects, whether treated or not and regardless of the assigned arm.*

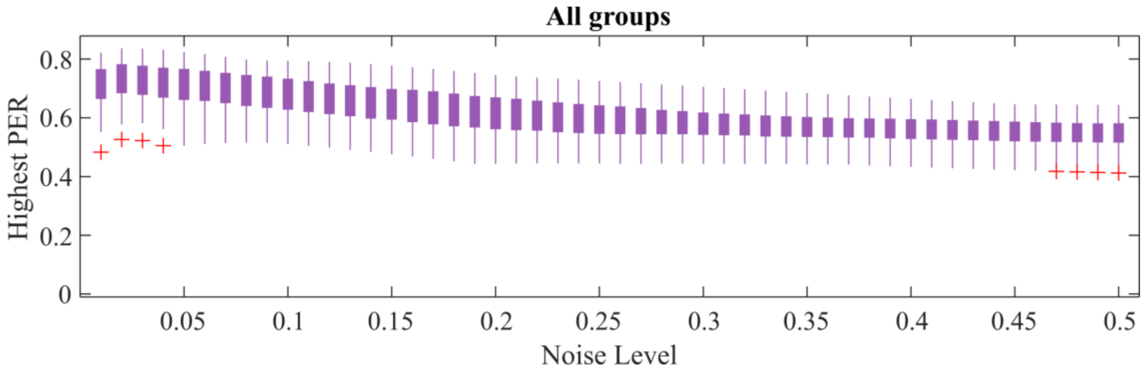


***Figure S5. Highest PER across noise levels****. The patient distribution of the highest perturbation effectivity to recovery (PER) values across areas is plotted as a function of the noise level. The distributions represent all subjects, whether treated or not and regardless of the assigned arm.*

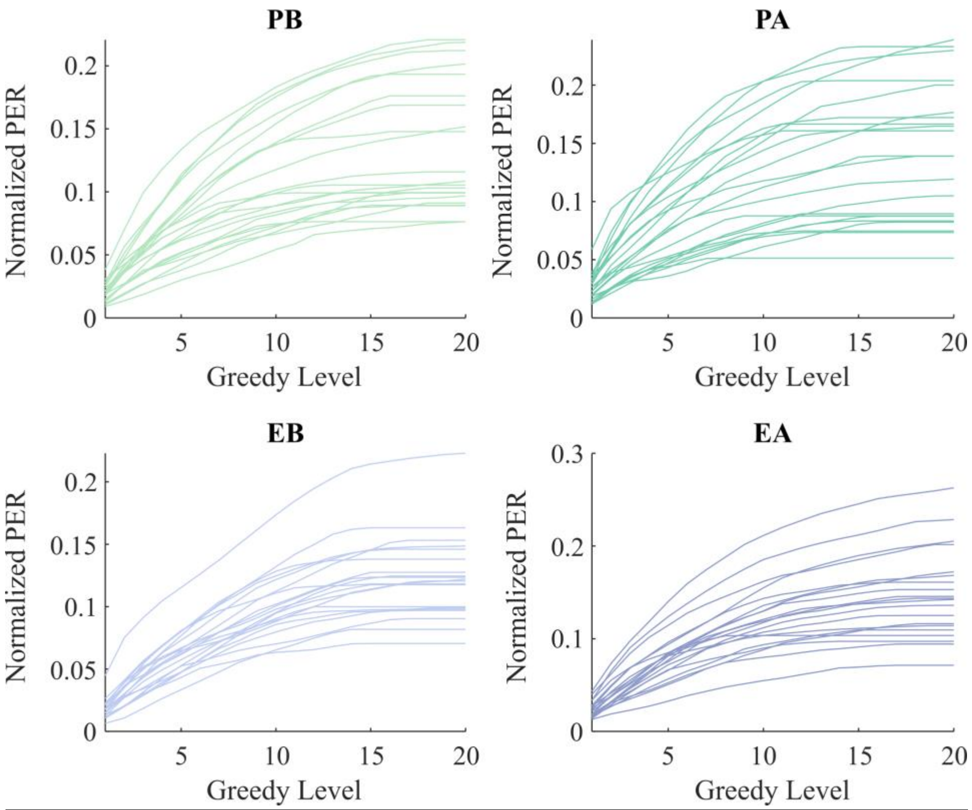


***Figure S6. Greedy multi-site perturbation trajectories. A)*** *The figure shows normalized PER trajectories for all PB (upper left), PA (upper right), EB (bottom left) and EA (bottom right) groups arising from PER maximization.*

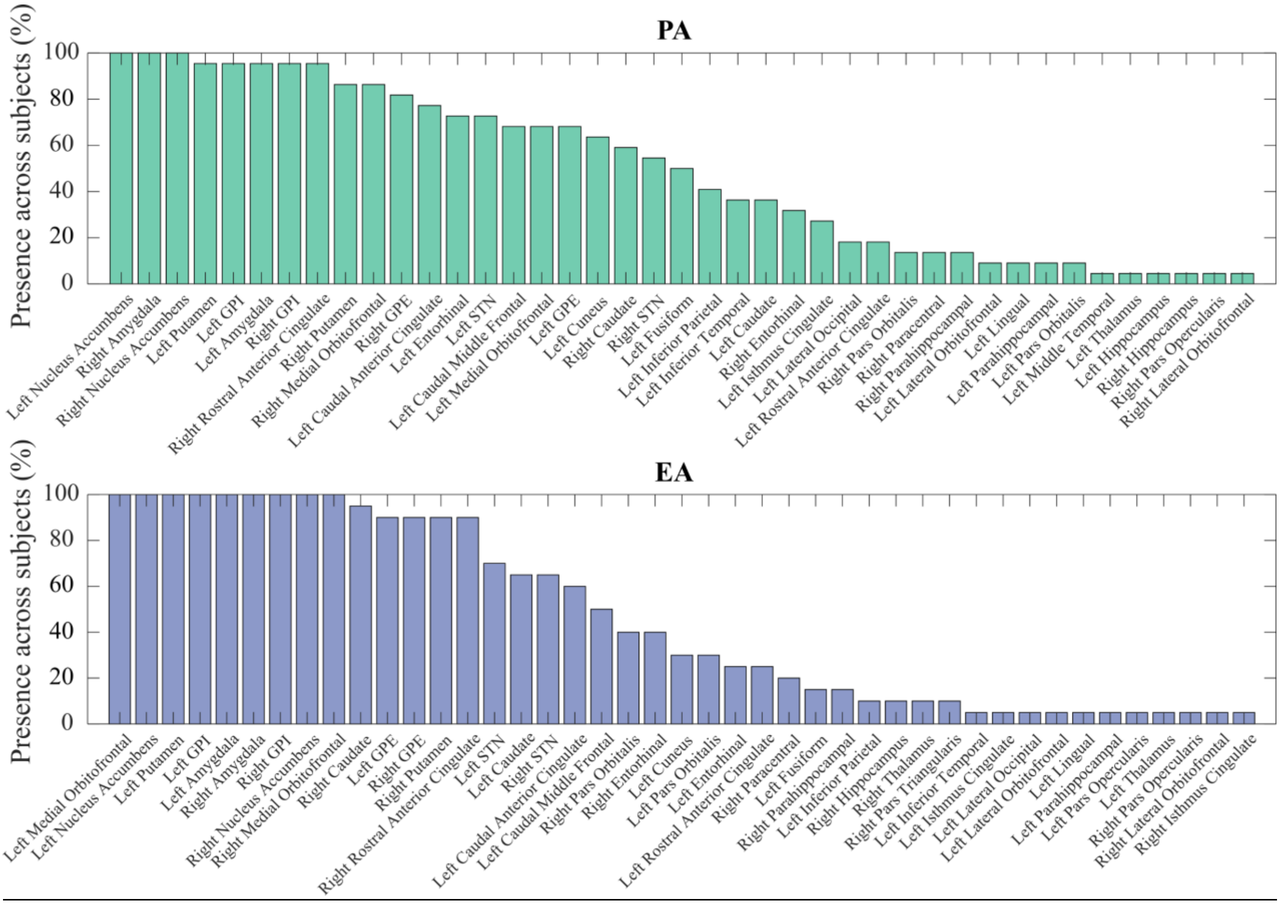


***Figure S7*. *Brain regions included in greedy multi-site perturbations.*** *The figure shows the presence of regions in the 20-area greedy multi-site perturbation strategy across patients, for both psilocybin and escitalopram arms after treatment (PA and EA, respectively).*
